## Supplementary information for "A Universal, Single-Component Multilayered Self-Assembling Protein Nanoparticle Vaccine Based on Extracellular Domains of Matrix Protein 2 Against Both Influenza A and B"

### Figure S1

a **Construct sequence**

**>M2exA3B3-5GS-1TD0 (A-A-A-B-B-B)**

DAMKRGLCCVLLLCGAVFVSPSQEIHARFRRGARSRSLLTEVETPIRNEWGCRCNDSSDGGGGSLLTEVETPTRNGWESKSSDSSDGGGGSLLTEVETPTRSEWESRSSGSSDGGGGML
EPFQIGGGGMLEPFQIGGGGMLEPFQIGGGGSEVRIFAGNDPAHTATGSSGISSPTPALTPLMLDEATGKLVVWDGQKAGSAVGILVLPLEGTETALTYYKSGTFATEAIHWPESVDEH
KKANAFAGSALSHAA

**>M2exA3B3-5GS-I3-01v9a (A-A-A-B-B-B)**

DAMKRGLCCVLLLCGAVFVSPSQEIHARFRRGARSRSLLTEVETPIRNEWGCRCNDSSDGGGGSLLTEVETPTRNGWESKSSDSSDGGGGSLLTEVETPTRSEWESRSSGSSDGGGGML
EPFQIGGGGMLEPFQIGGGGMLEPFQIASGGGGSAKLAEELQKKMEELFKKHKIVAVLRANSVEEAKEKALAVFEGGVHLIEITFTVPDADTVIKELSFLKEKGAIIGAGTVTSVEQCR
KAVESGAEFIVSPHLDAEITVFCLEKGVFYMPGVMTPTELVKAMKLGHNILKLFPGEVVGPQFVKAMKGPFPNVKFVPTGGVNLDNVCEWFKAGVLAVGVGSALVKGTPDEVREKAKAF
VEKIRGCTEGGGGSSPAVDIGDRLDELEKALEALSAEDGHDDVGQRLESLLRRWNSRRADGSAKFVAAWTLKAAA

**>M2exB3A3-5GS-I3-01v9a (B-B-B-A-A-A)**

DAMKRGLCCVLLLCGAVFVSPSQEIHARFRRGARSRMLEPFQIGGGGMLEPFQIGGGGMLEPFQIGGGGSLLTEVETPIRNEWGCRCNDSSDGGGGSLLTEVETPTRNGWESKSSDSSD
GGGGSLLTEVETPTRSEWESRSSGSSDASGGGGSAKLAEELQKKMEELFKKHKIVAVLRANSVEEAKEKALAVFEGGVHLIEITFTVPDADTVIKELSFLKEKGAIIGAGTVTSVEQCR
KAVESGAEFIVSPHLDAEITVFCLEKGVFYMPGVMTPTELVKAMKLGHNILKLFPGEVVGPQFVKAMKGPFPNVKFVPTGGVNLDNVCEWFKAGVLAVGVGSALVKGTPDEVREKAKAF
VEKIRGCTEGGGGSSPAVDIGDRLDELEKALEALSAEDGHDDVGQRLESLLRRWNSRRADGSAKFVAAWTLKAAA

**>M2e(AB)x3-5GS-I3-01v9a (A-B-A-B-A-B)**

DAMKRGLCCVLLLCGAVFVSPSQEIHARFRRGARSRSLLTEVETPIRNEWGCRCNDSSDGGGGMLEPFQIGGGGSLLTEVETPTRNGWESKSSDSSDGGGGMLEPFQIGGGGSLLTEVE
TPTRSEWESRSSGSSDGGGGMLEPFQIASGGGGSAKLAEELQKKMEELFKKHKIVAVLRANSVEEAKEKALAVFEGGVHLIEITFTVPDADTVIKELSFLKEKGAIIGAGTVTSVEQCR
KAVESGAEFIVSPHLDAEITVFCLEKGVFYMPGVMTPTELVKAMKLGHNILKLFPGEVVGPQFVKAMKGPFPNVKFVPTGGVNLDNVCEWFKAGVLAVGVGSALVKGTPDEVREKAKAF
VEKIRGCTEGGGGSSPAVDIGDRLDELEKALEALSAEDGHDDVGQRLESLLRRWNSRRADGSAKFVAAWTLKAAA

**>M2e(BA)x3-5GS-I3-01v9a (B-A-B-A-B-A)**

DAMKRGLCCVLLLCGAVFVSPSQEIHARFRRGARSRMLEPFQIGGGGSLLTEVETPIRNEWGCRCNDSSDGGGGMLEPFQIGGGGSLLTEVETPTRNGWESKSSDSSDGGGGMLEPFQI
GGGGSLLTEVETPTRSEWESRSSGSSDASGGGGSAKLAEELQKKMEELFKKHKIVAVLRANSVEEAKEKALAVFEGGVHLIEITFTVPDADTVIKELSFLKEKGAIIGAGTVTSVEQCR
KAVESGAEFIVSPHLDAEITVFCLEKGVFYMPGVMTPTELVKAMKLGHNILKLFPGEVVGPQFVKAMKGPFPNVKFVPTGGVNLDNVCEWFKAGVLAVGVGSALVKGTPDEVREKAKAF
VEKIRGCTEGGGGSSPAVDIGDRLDELEKALEALSAEDGHDDVGQRLESLLRRWNSRRADGSAKFVAAWTLKAAA

b **SDS Page**

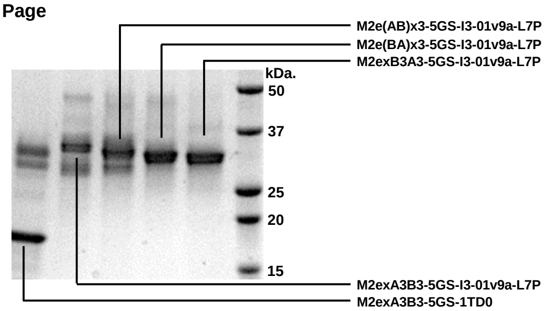


c **nsEM images**

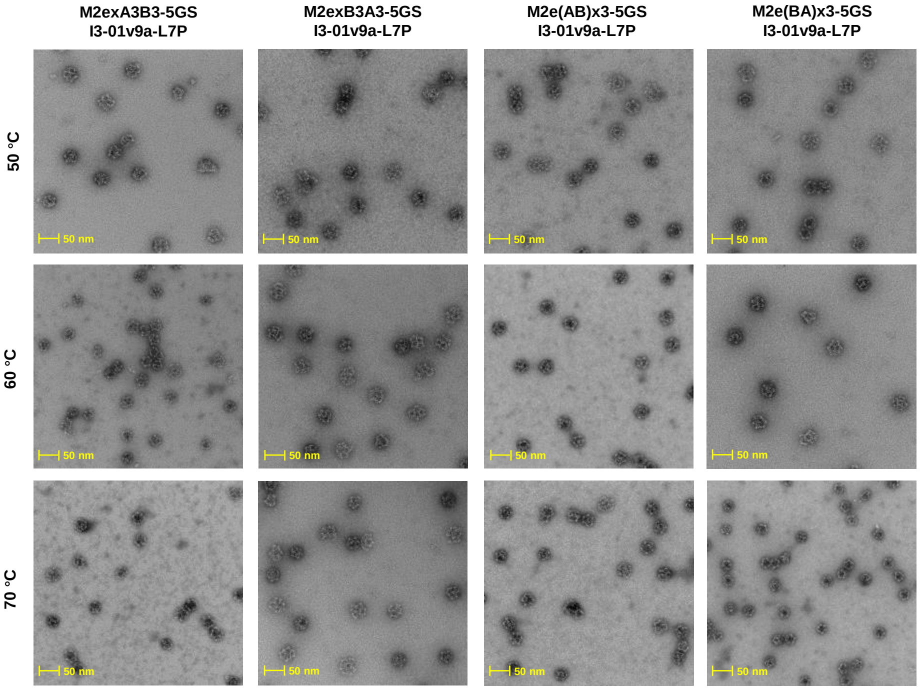

### Figure S1

d **ELISA profiles**

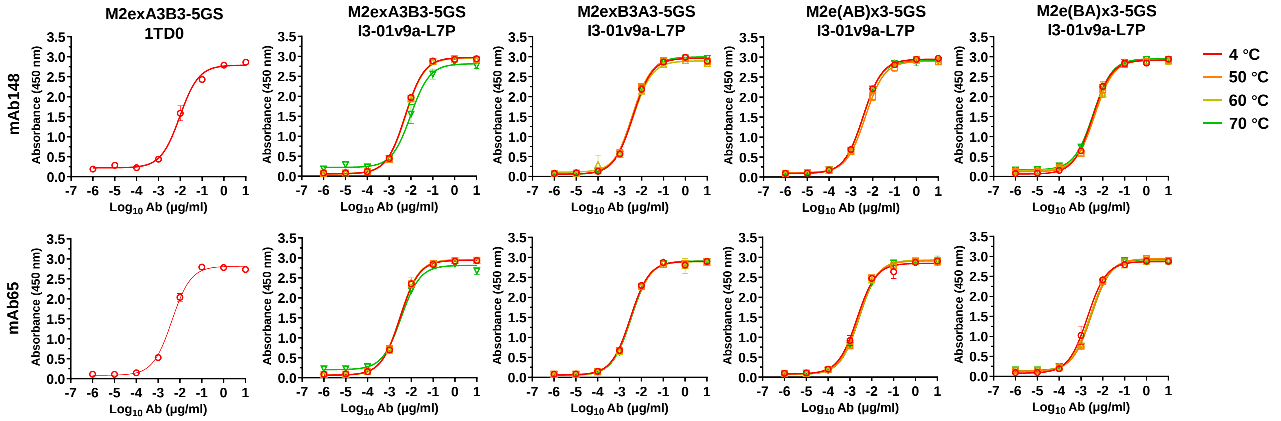


| $EC_{50}$ (ug/mL) | mAb148 | mAb65 |
|---|---|---|
| M2exA3B3-5GS-1TD0, 4 °C | 0.009586 | 0.004276 |
| M2exA3B3-5GS-I3-01v9a-L7P, 4 °C | 0.00548 | 0.00306 |
| M2exA3B3-5GS-I3-01v9a-L7P, 50 °C | 0.005598 | 0.003042 |
| M2exA3B3-5GS-I3-01v9a-L7P, 60 °C | 0.005624 | 0.003002 |
| M2exA3B3-5GS-I3-01v9a-L7P, 70 °C | 0.00976 | 0.003431 |
| M2exB3A3-5GS-I3-01v9a-L7P, 4 °C | 0.004018 | 0.003108 |
| M2exB3A3-5GS-I3-01v9a-L7P, 50 °C | 0.003661 | 0.003221 |
| M2exB3A3-5GS-I3-01v9a-L7P, 60 °C | 0.004132 | 0.003153 |
| M2exB3A3-5GS-I3-01v9a-L7P, 70 °C | 0.004012 | 0.003418 |
| M2e(AB)x3-5GS-I3-01v9a-L7P, 4 °C | 0.003651 | 0.002104 |
| M2e(AB)x3-5GS-I3-01v9a-L7P, 50 °C | 0.004584 | 0.002455 |
| M2e(AB)x3-5GS-I3-01v9a-L7P, 60 °C | 0.003747 | 0.002619 |
| M2e(AB)x3-5GS-I3-01v9a-L7P, 70 °C | 0.003616 | 0.002702 |
| M2e(BA)x3-5GS-I3-01v9a-L7P, 4 °C | 0.003397 | 0.001969 |
| M2e(BA)x3-5GS-I3-01v9a-L7P, 50 °C | 0.004113 | 0.002994 |
| M2e(BA)x3-5GS-I3-01v9a-L7P, 60 °C | 0.004305 | 0.002607 |
| M2e(BA)x3-5GS-I3-01v9a-L7P, 70 °C | 0.003598 | 0.003027 |

**Fig. S1. Design and in vitro characterization of influenza A-B M2e-presenting SApNPs.** (**a**) Construct sequences of influenza A-B M2e trimer and I3-01v9a-L7P SApNPs, with the gene fragments of sequence, restriction site, flexible linker, human, avian/swine, and human/swine matrix protein 2 extra-virion domain from influenza A virus (residue: 2-24), human matrix protein 2 extra-virion domain from influenza B virus (residue: 2-9), SApNP-forming subunit, trimerization domain (PDB: 1TD0), locking domain (LD), and PADRE highlighted in yellow, teal, green, gray, red, olive green, pink, cyan, and light blue, respectively. (**b**) SDS-PAGE analysis of various influenza A-B M2e trimer and SApNP constructs under reducing conditions. Each well was loaded with 2 μg of the appropriate protein. (**c**) Negative-stain EM images of influenza A-B M2e-presenting SApNPs after heat treatment at 50, 60, and 70 °C (**d**) ELISA showing influenza A-B M2e-based immunogen binding to mAb148 and mAb65.

### Figure S2

**a** **Mouse serum ELISA $EC_{50}$ titers**

| Antigen | EC50 titers (week 2) | | | | | | | | | | Geometric Mean |
|---|---|---|---|---|---|---|---|---|---|---|---|
| | M1 | M2 | M3 | M4 | M5 | M6 | M7 | M8 | M9 | M10 | |
| M2exA3B3 1TD0 trimer | 326.5 | 270.7 | 612.4 | 363.5 | 871 | 289.2 | 89.47 | 16.2 | 69.89 | 242.9 | 203.5 |
| M2exA3B3 I3-01v9a SApNPs | 2460 | 1535 | 1769 | 2592 | 3088 | 3459 | 1516 | 3932 | 2103 | 1771 | 2297.9 |
| M2exB3A3 I3-01v9a SApNPs | 2458 | 4069 | 2739 | 3447 | 4099 | 6917 | 2441 | 4928 | 2460 | 2112 | 3329.3 |
| M2e(AB)x3 I3-01v9a SApNPs | 1076 | 473.6 | 1660 | 1356 | 1792 | 2069 | 1252 | 1925 | 1044 | 1816 | 1345.4 |
| M2e(BA)x3 I3-01v9a SApNPs | 1484 | 968 | 1865 | 1444 | 1581 | 1410 | 4345 | 2430 | 1539 | 1219 | 1672.1 |

| Antigen | EC50 titers (week 5) | | | | | | | | | | Geometric Mean |
|---|---|---|---|---|---|---|---|---|---|---|---|
| | M1 | M2 | M3 | M4 | M5 | M6 | M7 | M8 | M9 | M10 | |
| M2exA3B3 1TD0 trimer | 174152 | 13817 | 74732 | 20320 | 22758 | 8525 | 16548 | 5408 | 7568 | 8970 | 18340.7 |
| M2exA3B3 I3-01v9a SApNPs | 210684 | 180258 | 169830 | 199148 | 245451 | 203008 | 222892 | 193213 | 209258 | 83127 | 185385.1 |
| M2exB3A3 I3-01v9a SApNPs | 109611 | 83460 | 179999 | 222131 | 120769 | 102311 | 151243 | 177892 | 223975 | 117397 | 141409.2 |
| M2e(AB)x3 I3-01v9a SApNPs | 44747 | 90241 | 67911 | 122343 | 94670 | 130642 | 57744 | 110003 | 160877 | 114523 | 93031.1 |
| M2e(BA)x3 I3-01v9a SApNPs | 84970 | 42554 | 70252 | 79832 | 80588 | 113303 | 121094 | 84294 | 100677 | 74586 | 82263.3 |

**b** **Statistical analysis**

**Week 2**

| One-way ANOVA with Tukey's multiple comparisons test (w2) | Statistics | Adjusted P Value |
|---|---|---|
| M2exA3B3 1TD0 trimer-w2 vs. M2exA3B3 I3-01v9a SApNPs-w2 | **** | <0.0001 |
| M2exA3B3 1TD0 trimer-w2 vs. M2exB3A3 I3-01v9a SApNPs-w2 | **** | <0.0001 |
| M2exA3B3 1TD0 trimer-w2 vs. M2e(AB)x3 I3-01v9a SApNPs-w2 | ns | 0.0604 |
| M2exA3B3 1TD0 trimer-w2 vs. M2e(BA)x3 I3-01v9a SApNPs-w2 | ** | 0.0051 |
| M2exA3B3 I3-01v9a SApNPs-w2 vs. M2exB3A3 I3-01v9a SApNPs-w2 | ns | 0.056 |
| M2exA3B3 I3-01v9a SApNPs-w2 vs. M2e(AB)x3 I3-01v9a SApNPs-w2 | ns | 0.1387 |
| M2exA3B3 I3-01v9a SApNPs-w2 vs. M2e(BA)x3 I3-01v9a SApNPs-w2 | ns | 0.5993 |
| M2exB3A3 I3-01v9a SApNPs-w2 vs. M2e(AB)x3 I3-01v9a SApNPs-w2 | **** | <0.0001 |
| M2exB3A3 I3-01v9a SApNPs-w2 vs. M2e(BA)x3 I3-01v9a SApNPs-w2 | *** | 0.001 |
| M2e(AB)x3 I3-01v9a SApNPs-w2 vs. M2e(BA)x3 I3-01v9a SApNPs-w2 | ns | 0.8826 |

**Week 5**

| One-way ANOVA with Tukey's multiple comparisons test (w5) | Statistics | Adjusted P Value |
|---|---|---|
| M2exA3B3 1TD0 trimer-w5 vs. M2exA3B3 I3-01v9a SApNPs-w5 | **** | <0.0001 |
| M2exA3B3 1TD0 trimer-w5 vs. M2exB3A3 I3-01v9a SApNPs-w5 | **** | <0.0001 |
| M2exA3B3 1TD0 trimer-w5 vs. M2e(AB)x3 I3-01v9a SApNPs-w5 | * | 0.0124 |
| M2exA3B3 1TD0 trimer-w5 vs. M2e(BA)x3 I3-01v9a SApNPs-w5 | ns | 0.0808 |
| M2exA3B3 I3-01v9a SApNPs-w5 vs. M2exB3A3 I3-01v9a SApNPs-w5 | ns | 0.1773 |
| M2exA3B3 I3-01v9a SApNPs-w5 vs. M2e(AB)x3 I3-01v9a SApNPs-w5 | *** | 0.0001 |
| M2exA3B3 I3-01v9a SApNPs-w5 vs. M2e(BA)x3 I3-01v9a SApNPs-w5 | **** | <0.0001 |
| M2exB3A3 I3-01v9a SApNPs-w5 vs. M2e(AB)x3 I3-01v9a SApNPs-w5 | ns | 0.0849 |
| M2exB3A3 I3-01v9a SApNPs-w5 vs. M2e(BA)x3 I3-01v9a SApNPs-w5 | * | 0.0132 |
| M2e(AB)x3 I3-01v9a SApNPs-w5 vs. M2e(BA)x3 I3-01v9a SApNPs-w5 | ns | 0.944 |

**Fig. S2. Immunogenicity of influenza A-B M2e-based vaccines in mice.** (**a**) $EC_{50}$ titers for M2exA3B3 trimer and I3-01v9a SApNP vaccine-immune sera binding to sequence-matched M2e coating antigen (M2exA3B3-5GS-foldon, M2exB3A3-5GS-foldon, M2e(AB)x3-5GS-foldon, and M2e(BA)x3-5GS-foldon). Color coding indicates the magnitude of $EC_{50}$ titers (green to red: low to high). (**b**) Table of statistical analysis was performed using a one-way ANOVA followed by Tukey's multiple-comparison *post hoc* test for each timepoint. $EC_{50}$ titers were calculated in GraphPad Prism 10.2.3. For significance, ns (not significant), *$p < 0.05$, **$p < 0.01$, ***$p < 0.001$, and ****$p < 0.0001$.
